# Embryogenesis, polyembryony, and settlement in the gorgonian *Plexaura homomalla*

**DOI:** 10.1101/2020.03.19.999300

**Authors:** Christopher D. Wells, Kaitlyn J. Tonra, Howard R. Lasker

## Abstract

Understanding the ontogeny and reproductive biology of reef-building organisms can shed light on patterns of population biology and community structure. This knowledge is particularly important for Caribbean octocorals, which seem to be more resilient to long-term environmental change than scleractinian corals and provide some of the same ecological services. We monitored the development of the black sea rod *Plexaura homomalla*, a common, widely distributed octocoral on shallow Caribbean reefs, from eggs to 3-polyp colonies over the course of 73 days. In aquaria on St John, U.S. Virgin Islands, gametes were released in spawning events three to six days after the July full moon. Cleavage started 3 hours after fertilization and was holoblastic, equal, and radial. Embryos were positively buoyant until becoming planulae. Planulae were competent after 4 days. Symbiodiniaceae began infecting polyps at around 8 days post fertilization. Development was typical for Caribbean octocorals, except for the occurrence of a novel form of asexual reproduction in octocorals: polyembryony. Fragmentation of embryos during development may represent a temporally varied tradeoff between number and size of propagules, in which large eggs have higher fertilization rates followed by polyembryony, which maximizes the number of surviving recruits by generating more, albeit smaller, larvae. Polyembryony may contribute to the success of some gorgonians on Caribbean reefs as other anthozoans are in decline.

## 1. INTRODUCTION

Understanding the ontogeny, reproductive biology, and embryonic investment of reef-building organisms can shed light on large-scale patterns of population biology and community structure (Babcock et al., 1986; Baums et al., 2006; Baird et al., 2009). For example, this information can help explain the resilience of ecosystems and populations to disturbance (Nyström and Folke, 2001; Waples et al., 2009). This is particularly important for taxa like Caribbean octocorals, which provide some of the same ecosystem services as scleractinian corals, such as three-dimensional structure for fish grazing (Tsounis et al., 2016) and the promotion of coral recruitment (Privitera-Johnson et al., 2015). While scleractinian cover on Caribbean reefs has declined (Hughes, 1994; Gardner et al., 2003), octocoral abundance has been increasing (Ruzicka et al., 2013; Lenz et al., 2015; Sánchez et al., 2019).

Octocorals seem to be more resilient to long-term disturbance than scleractinian corals, possibly due to increased recruitment success (Edmunds and Lasker, 2016; Tsounis and Edmunds, 2017; Lasker et al., 2020). Successful recruitment is necessary for the maintenance of a population, and characteristics of reproduction and development in octocorals may help explain their success in Caribbean reefs. The general patterns of sexual and asexual reproduction in octocorals are well-documented. Species are primarily gonochoric and either broadcast-spawn gametes or brood embryos (Kahng et al., 2011). Eggs are frequently yolky, cleavage is superficial and holoblastic, and the blastula is solid (Lasker and Kim, 1996; Dahan and Benayahu, 1998). Despite our in-depth knowledge of oogenesis and reproductive habits, embryogenesis and early development in octocorals are understudied and the ontogenies of many conspicuous species have yet to be studied in detail.

In this study, we monitored the previously undescribed development of the black sea rod *Plexaura homomalla* (Esper, 1794), a common, widely-distributed, fast-growing, gonochoric, broadcast-spawning octocoral found in shallow Caribbean reefs (reviewed in Kim and Lasker, 1997). This work provides a detailed timeline of early development and introduces a phenomenon, polyembryony (*sensu* Allen et al., 2018), which had not previously been reported in octocorals but has been described in scleractinians (Heyward and Negri, 2012; Chamberland et al., 2017) and many other phyla (reviewed in Craig et al., 1997). Fragmentation of embryos during development may represent a tradeoff between survival of individuals and survival of the clone, and could play a key role in recruitment of some gorgonians and their subsequent success on Caribbean reefs.

## 2. METHODS

Female and male branches of *Plexaura homomalla* were collected from an octocoral-dominated reef off of St. John, USVI on July 14-15, 2019 (18.345 °N, 64.681 °W) between 3.0 and 5.0 m depth. Colonies were determined to be sexually mature in the field. Colonies were transported to the Virgin Islands Environmental Resource Station and maintained in a sea table with unfiltered running seawater. Exchange rate in the sea table was approximately 200 L/h (60% of the volume hourly). Two EcoPlus 1/10 HP water chillers (Hawthorne Gardening Company, Vancouver, WA) were run in series, set to a temperature of 27°C to reduce temperature variability. This reduced water temperature 0.5°C during the day (daily temperature range: 27.0-29.4°C) and maintained temperature at 27°C at night. Two submersible circulation pumps (Maxi-Jet 900 and Mini-Jet 404 [Marineland Spectrum Brands Pet LLC, Blacksburg, VA]) were placed within the tank to create a circular flow within the sea table. Flow rate was high enough within the sea table to cause the branch tips of the colonies to move slightly when an eddy caught them, but they were not consistently shaking.

The sea tables were observed for signs of spawning (e.g., eggs on the surface or in the water column) from July 17th, 2019, one day after the full moon, until July 23rd, 2019. At the start of spawning, pumps and water input were shut off. Gametes were collected with 60-mL syringes as they were released from the colonies and were stored in water from the tank for 30 minutes to ensure fertilization. The time at which gametes were mixed is referred to as t = 0 because we were unable to determine the exact moment of fertilization for any given egg.

Embryo development was observed over two nights (July 20 and 21). On the first night, 100 oocytes were collected and the number of cells was counted every half-hour from 2.5 hours post fertilization (hpf) to 7 hpf and then hourly until 10 hpf. Cells were counted until the embryos had 64 cells. Gross morphology observations were made at 15, 18, 21, and 24 hpf, and then daily for three days. Planulae were allowed to settle on 14 x 14 x 1 cm stoneware clay tiles where they metamorphosed into polyps. The tiles were deployed onto an octocoral-dominated reef (18.309 °N, 64.719 °W) and the number of polyps in each colony was counted at 14, 33, and 71 days post fertilization. On the second night of spawning, a more detailed study of early development was conducted by counting the number of cells in 11 embryos every ten minutes for the first two hours and then every five minutes for the next four hours. We stopped counting the number of cells after an individual reached 128 cells as further divisions became difficult to detect accurately.

## 3. RESULTS

Approximately two hours after sunset on the evenings of July 19-22, 2019, 3-6 days after the full moon, *Plexaura homomalla* spawned in the tanks. Spawning was synchronous, but females were observed spawning first. Pumps were not turned off until the first eggs were observed, so early release of sperm may have been missed because sperm are released in indistinct clouds. For each colony, gametes were released for two hours on up to three consecutive nights, but total spawning lasted four hours each night for four consecutive nights. Only colonies with extended polyps released gametes. Total number of eggs collected on July 19th, 20th, 21st, and 22nd were 35, 7381, 6002, and 123, respectively, from 10 female 15-cm branches (approximately 1350 eggs/branch). All eggs that were released were collected on July 19th and 22nd and nearly all were collected on July 20th and 21st.

Eggs and early-stage embryos were positively buoyant and some became trapped in the surface tension. Eggs were spherical and 600-800 µm in diameter with heavy investment in yolk (Fig. 1A). During the first night of observations, 15 of 100 eggs died in the first three hours of our monitoring, possibly due to a lack of fertilization; this suggests a fertilization rate of at least 85%. Cleavage began 3 hpf and was holoblastic, equal, and radial (Fig. 1B-C, Tab. 1). Cleavage started at one pole of the oocyte and traveled across with increased furrow formation after each division. Cleavages to 8, 16, 32, and 64-celled embryos led to blastulae with distinct blastomeres (Fig. 1B-C). Division from 8 cells to 64 cells took approximately 45 mins (Fig. 2, Tab. 1). Embryos that did not progress through stages at a steady rate (i.e., too fast or slow) frequently died. A spherical stereoblastula was formed 4.9 hpf and by 15 hpf all embryos were irregularly shaped gastrula shaped similarly to a raisin (Fig. 1D, Tab. 1).

**Table 1.** Median and range of time to reach stages of development in the gorgonian *Plexaura homomalla.*

| Development Stage | Median Time | Range |
| --- | --- | --- |
| Oocyte | 0 hours | 0 – 3.7 hours |
| 2-cell | 3.0 hours | 2.8 – 3.7 hours |
| 4-cell | 3.3 hours | 2.8 – 3.8 hours |
| 8-cell | 3.4 hours | 2.8 – 3.8 hours |
| 16-cell | 3.7 hours | 3.0 – 3.9 hours |
| 32-cell | 3.8 hours | 3.4 – 4.2 hours |
| 64-cell | 4.3 hours | 4.1 – 4.6 hours |
| Stereoblastula | 4.9 hours | 4.8 – 6 hours |
| Gastrula | 15 hours | 5 – 36 hours |
| Planula | 3 days | 2.5 – 4 days |
| Settlement (polyp) | 5 days | 5 – 8 days |
| 3-polyp colony | 71 days |  |

**Figure 1.**
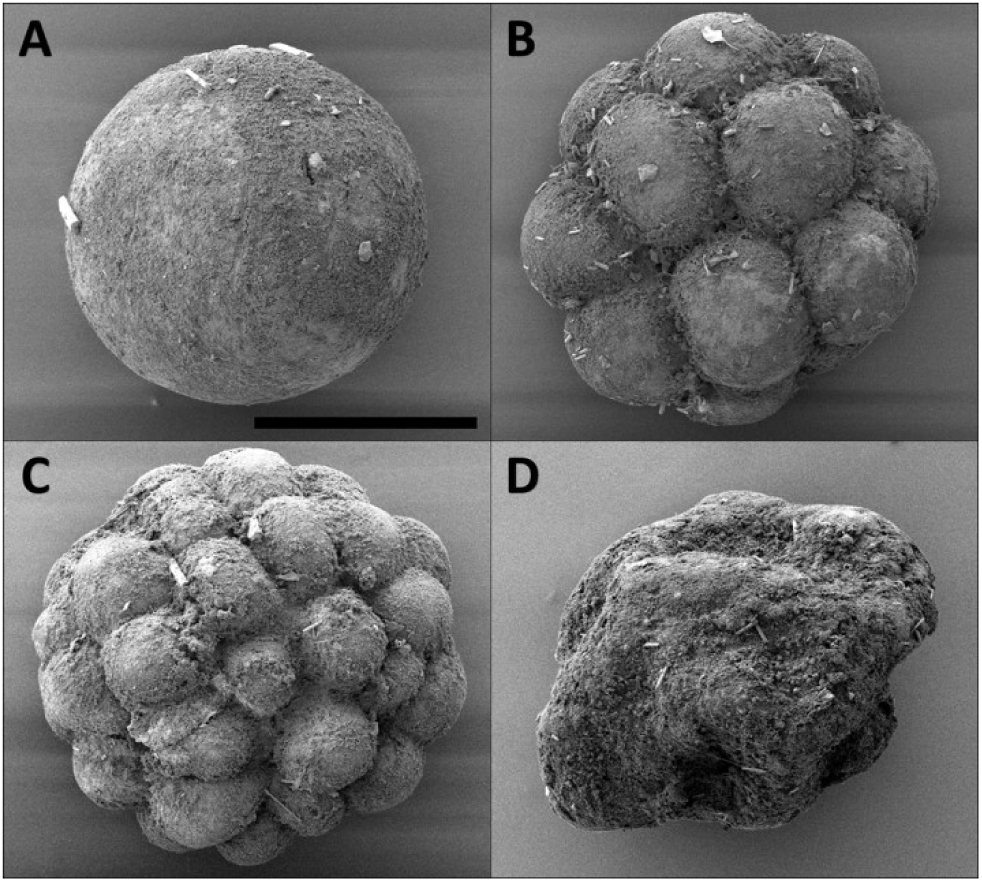
Scanning electron microscope images of early development events of the gorgonian *Plexaura homomalla*: (A) egg/embryo (scale bar A-D = 300 μm) (B) 16-cell embryo, (C) 32-cell embryo, and (D) early-stage gastrula.

**Figure 2.**
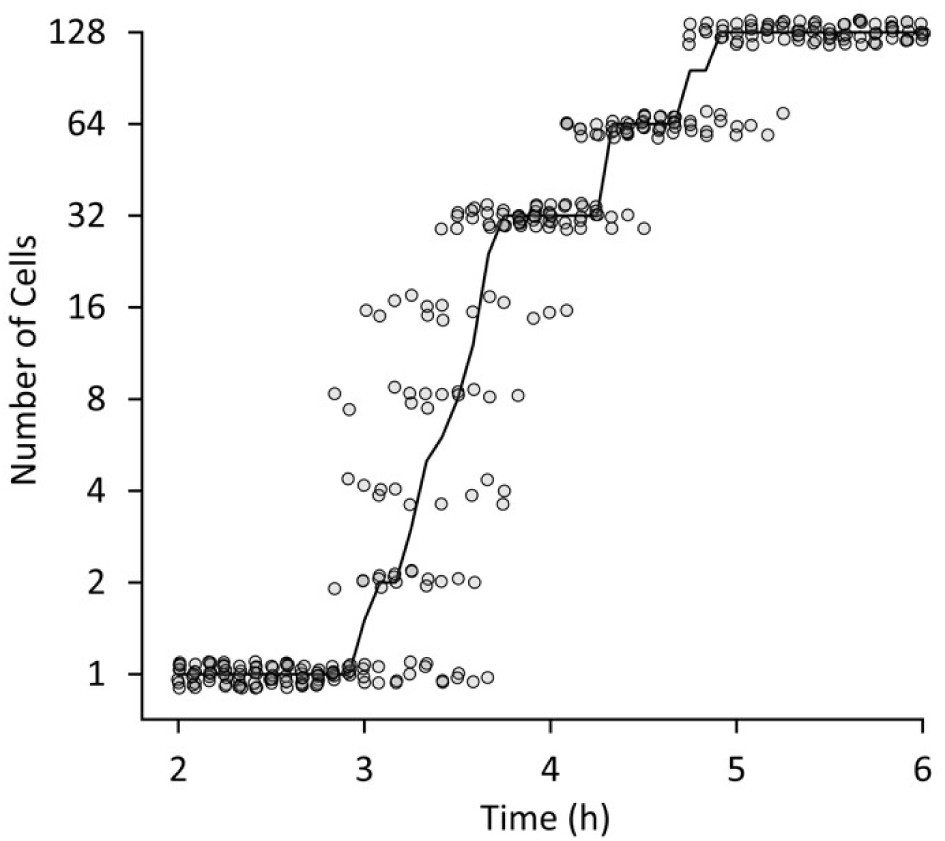
Development of the gorgonian *Plexaura homomalla* from two to six hours after fertilization, checked at 5-minute intervals. Each point represents one observation. Cells start dividing around 3 hours after fertilization and had all reached 128+ cells by 5.5 hours after fertilization. The solid black line is the median number of cells. Values have been spread on the x- and y-axis to reduce overlap.

During the second day, embryos became ovoid and were less positively buoyant and by 3 days, embryogenesis was complete and planulae had formed (Fig 3A, Tab. 1). Planulae were slightly negatively buoyant, pear-shaped to ovoid, and were able to move at approximately 0.5 cm/s. They had the ability to become elongated but could quickly return to a pear-shape. Planulae could be found crawling on the surfaces of the culture container, but did not stick inside the glass pipettes during water changes. By day four, planulae were clearly competent to settle. They readily attached to surfaces at that time and it was difficult to transfer planulae without forceful flushing of the water to remove them from the sides of the pipette.

**Figure 3.**
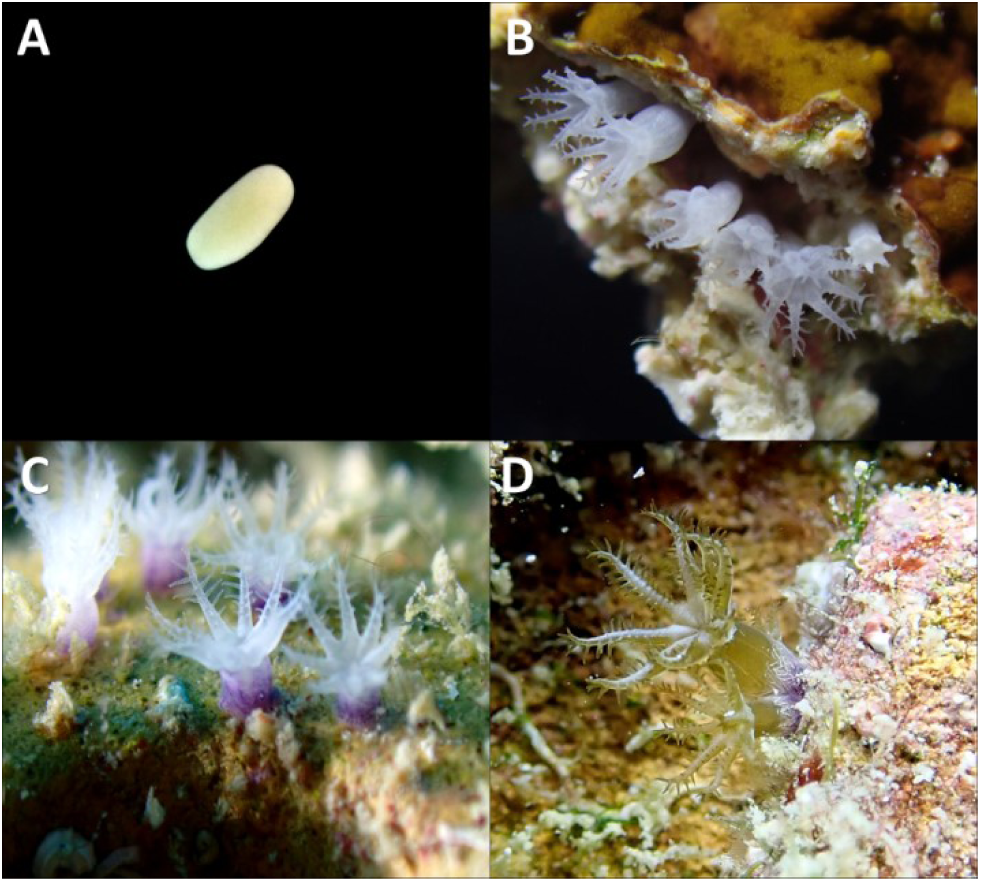
Photographs of the early development of the gorgonian *Plexaura homomalla*: (A) planula larva (approximately 1 mm long), (B) 12 days old polyps (polyp columns are approximately 0.1 cm tall for B-C) (C) 18 days old polyps with developed purple sclerites and (D) 71 days old colonies (column of the main polyp of the colony is approximately 0.4 cm tall).

Planulae began settling 5 days after fertilization. Settlement was asynchronous; some planulae settled as late as 8 days after fertilization. Typically, planulae landed on a surface, crawled around, and either attached to the substratum or swam back into the water column. Endosymbiotic dinoflagellates (Symbiodiniaceae) were not apparent at settlement (Fig. 3B), but tentacles on some polyps had begun to turn brown by 8 days after fertilization. Between days 12 and 18, purple sclerites were being produced (Fig. 3C). Colonies ranging from two to four polyps formed between 33 and 71 days after fertilization (Fig. 3D, Tab. 1).

### 3.1 Polyembryony

After reaching 64 cells, some embryos fragmented and led to an increase in the number of embryos (Fig. 4). While we did not directly observe the fragmentation process, the increase in the number and the presence of smaller than normal embryos (i.e., approximately half the size) strongly suggests polyembryony. We were not able to monitor developmental success of individual fragmented embryos, but the presence and then disappearance of extremely small embryos suggests unequal fragmenting occurred but frequently resulted in the death of the smaller half.

**Figure 4.**
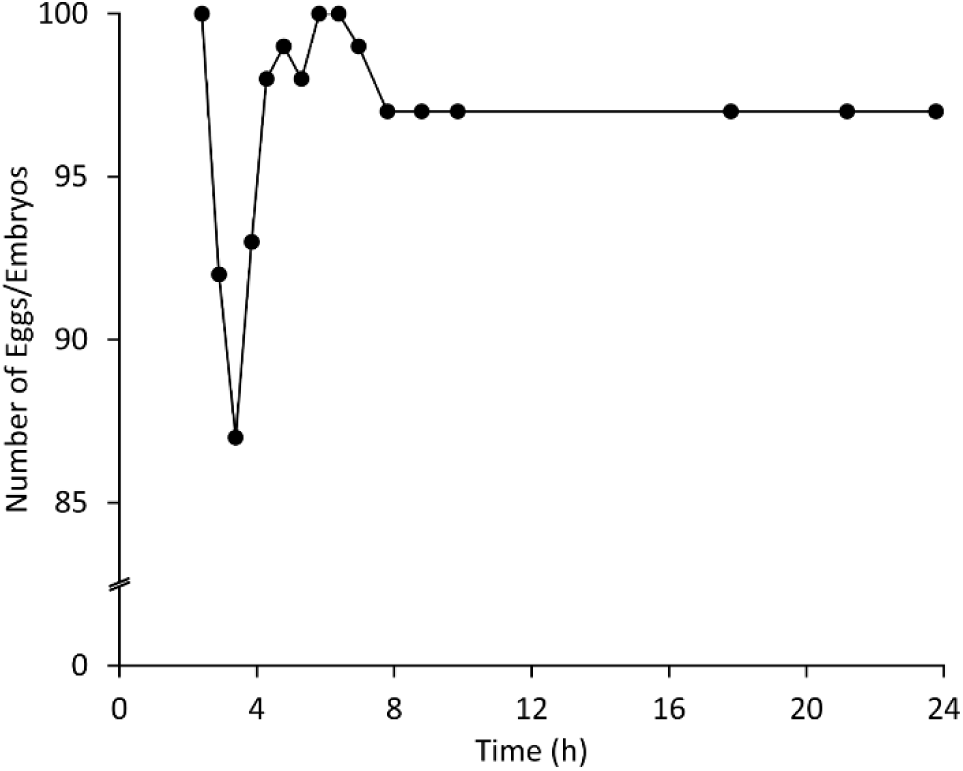
Number of eggs/embryos of the gorgonian *Plexaura homomalla* alive in the first 24 hours of development starting with 100 eggs. Mortality was high between hours 2.5 and 3.5. At 3.5 hours, some 64+ cell embryos broke into fragments. The number of embryos stabilized after eight hours during the blastula stage.

## 4. DISCUSSION

This study provides the first detailed description of the spawning and embryogenesis of *Plexaura homomalla* (Fig. 1-3, Tab. 1). Compared to field observations from Venezuela (Bastidas et al., 2005), we observed spawning 1-2 lunar cycles earlier, spawning lasted for twice the duration on each night, and took place over twice as many nights. Colonies released few gametes on the first and last days of spawning, so observations in a laboratory setting may be advantageous for detecting early spawners. The temporal difference between spawning times at different sites has been observed with other octocorals and has been related to differences in local temperature (Pakes and Woollacott, 2008; De Putron and Ryland, 2009). Alternatively, these differences could be an artefact of collecting and spawning pieces of colonies in a laboratory (e.g., light pollution or stress).

Development was typical for Caribbean octocorals (Benayahu, 1989; Dahan and Benayahu, 1998; Mandelberg-Aharon and Benayahu, 2015), except for the occurrence of polyembryony (Fig. 4). Polyembryony has been reported in reef-building scleractinian corals (Heyward and Negri, 2012; Chamberland et al., 2017), but never in octocorals. In both scleractinian studies reporting polyembryony, embryos generated from fragmentation developed into competent, albeit smaller, planulae that successfully settled. In the case of *P. homomalla*, we have inferred polyembryony from the increase in numbers of embryos and the presence of smaller than normal embryos in our cultures. Smaller embryos were approximately half the size of normal embryos, suggesting that fragmentation produced two clones from one embryo. However, we cannot exclude the possibilities that fragmentation could produce more than two clones or that one embryo can undergo fragmentation multiple times. Heyward and Negri (2012) found that during turbulent conditions, small breaking waves could induce fragmentation in scleractinians during the 2- to 16-cell stages. *P. homomalla* embryos were left in still water during this study and fragmented at a later stage (64+ cells), indicating that polyembryony in this taxon may be induced by a different trigger. Polyembryony has been induced in sea urchins exposed to low salinity environments, possibly due to reduced levels of Ca^2+^ (Allen et al., 2015), which is necessary for normal interactions between blastomeres (Vacquier and Mazia, 1968a, 1968b). While the trigger remains unknown, the result is the same - clonemates are produced from one fertilization event.

In the context of the advantages of sexual and asexual reproduction (Williams, 1975; Maynard Smith, 1978), polyembryony has been discussed as a fitness paradox (e.g., Ryland, 1996; Craig et al., 1997; Hughes et al., 2005) – it seemingly misses out on the full benefits of potentially generating more fit genotypes through sexual reproduction and the advantage of replicating the maternal genotype, a proven successful genotype, through asexual reproduction. Through polyembryony, at least two copies of each genotype are generated. Using Williams’ (1975) metaphor, identical “lottery tickets” are produced. This seemingly wasteful strategy has value if the probability of successful recruitment is extremely low and mostly independent of genotype. In this case, almost all lottery tickets are lost and having replicate tickets is no less advantageous than having twice as many different tickets. The probability of any one genotype surviving is far greater when there are two copies. If the loss of genetic diversity is low, then the success of polyembryony is likely to be based on the traditional tradeoff between the size and number of zygotes produced (*cf*. Stearns, 1977; Strathmann, 1977). In the case of *P. homomalla*, polyembryony provides the additional capability of changing the size to number relationship during the course of development.

Craig et al. (1997) suggested polyembryony may be favorable when the mother is unable to detect the ecological circumstances that the offspring will experience. Many animals that have exhibited polyembryony have long oogenesis times or extended larval development. For example, cyclostome bryozoans have a prolonged oogenic cycle. Their food source is temporally patchy (Ryland, 1996), but polyembryony can increase clutch size for this taxon in times of abundant food. *P. homomalla* also has a protracted oogenesis cycle (12 months; Fitzsimmons-Sosa et al., 2004). However, more eggs are present early in the oogenic cycle than fully develop (HRL, unpublished observations of fecundity in polyps one and two months prior to spawning), which suggests that unlike cyclostome bryozoans, *P. homomalla* may be able to respond to environmental cues during oogenesis and adjust the number of eggs produced. Furthermore, *P. homomalla* has lecithotrophic (i.e., non-feeding) larvae, making success of the larvae independent of planktonic food availability.

Polyembryony might also be favored when fertilization is constrained in some manner (Craig et al., 1997). Fertilization can be limited by phylogenetic constraints on egg production (Prodöhl et al., 1996), predation of eggs before fertilization (Chamberland et al., 2017), or by sperm availability (Lasker and Stewart, 1993). While many studies have demonstrated that sperm-limitation is possible in marine environments (e.g., Lasker and Stewart, 1993; Levitan and Petersen, 1995), fertilization success is generally high in octocorals (Coma and Lasker, 1997a, 1997b; Lasker, 2006). Producing large eggs increases the target area for sperm and improves fertilization success (Levitan, 1993). Polyembryony allows *P. homomalla* to obtain the benefits of having large eggs at the time of fertilization, and then have more numerous but smaller embryos and larvae over the remainder of development. In this case, polyembryony may allow an optimization of size and number of eggs that can be altered over the course of development. A test of this hypothesis will require a quantification of the relationship between egg size and fertilization success and the relationship between embryo size and successful recruitment.

While we know coral embryos can be triggered to fragment by physical disturbances such as waves (Heyward and Negri, 2012), other potential triggers such as temperature and salinity are not well understood. Inducing polyembryony may help in coral restoration efforts (Randall et al., 2020) or in experiments where the number of embryos is limited. Early development in octocoral taxa share many early ontogenetic features suggesting that these components have a broad phylogenetic basis. A comprehensive survey for polyembryony in octocorals is needed to determine whether this is a widespread phenomenon and important to the success of the taxon or limited to a smaller number of taxa and circumstances.

## ACKNOWLEDGEMENTS

This work was completed under permits from the Virgin Islands Division of Fish and Wildlife (DFW19010) and funded by the National Science Foundation (OCE 17-56381). We thank the Virgin Islands Environmental Resource Station (VIERS) and the University of the Virgin Islands (UVI) for laboratory space, A. Martinez-Quintana for field and laboratory assistance, and E.R. Anderson and M.A. Coffroth for comments on the manuscript.

## Notes

#### Summary of Updates

Addressing formatting issues (overlapping text and off-margins).

